# SNX-2112 inhibits the ERBB3^high^ tumor population in Plexiform Neurofibromas

**DOI:** 10.1101/2025.04.26.650740

**Authors:** Sajjad Khan, Oluwatosin Aina, Ximei Veneklasen, Benjamin Bruckert, Kimani Njoya, Donia Alson, Li Sun, Huda Zayed, Daochun Sun

**Author notes:** Correspondence, Address: 8701 W Watertown Plank Rd, Milwaukee, WI 53226, USA. Concordia University of Wisconsin, 12800 N Lake Shore Dr, Mequon, WI, 53080.

## Abstract

Plexiform neurofibroma (PN) is a monogenic disorder affecting the Schwann cell lineage in approximately half of Neurofibromatosis type 1 patients. It often manifests in early childhood and carries a risk of malignant transformation at puberty. While MEK inhibitors have recently been approved for pediatric and adult patients, their long-term clinical benefits remain uncertain due to adverse effects, particularly in pediatrics. The ERBB3^high^ stem-like tumor cell subpopulation reflects the intrinsic PN heterogeneity and can serve as alternative druggable targets. Characterization using molecular signatures also reveals the heterogeneity within the PN cell lines, and the PN drug screen analysis proposes that SNX-2112, an HSP90 inhibitor, can inhibit PN cell viability by targeting ERBB3-associated signaling. Our *in vitro* results demonstrated a significant viability reduction in ERBB3^high^ PN cells compared to the controls. Treatment with SNX-2112 led to a decrease in total ERBB3, p-ERBB3 (Tyr1289), and the downstream total AKT and p-AKT (Ser473) levels. Furthermore, the significant *in vivo* inhibitory efficacy of SNX-2112 was revealed using a transplantation model from mouse PN with *DhhCreNF1*^*f/f*^;*Luc* configuration. These findings support the efficacy of SNX-2112 as a potential alternative therapeutic strategy, highlighting its possible application in PN patients.

## Introduction

Plexiform neurofibromas (PNs) are challenging benign peripheral nerve sheath tumors, particularly associated with Neurofibromatosis type 1 (NF1). PNs have a high risk of progressing to malignant peripheral nerve sheath tumors (MPNSTs), which leads to the major mortality of NF1 patients. PNs arise from neural crest stem cells (NCSCs), and loss of function of the *NF1* gene in early Schwann cell lineage (SCL) contributes to tumorigenesis [1]. MEK inhibitors, selumetinib and mirdametinib, have been approved by the FDA to inhibit the downstream signaling of hyperactive Ras signaling in pediatric and adult patients [2, 3]. However, long-term adverse effects of MEK inhibitors have been generally reported in cancer patients, especially in children [4, 5]. PN has complex tumor intrinsic heterogeneity, and it is comprised of a significant population of stem-like tumor cells committed to Schwann cell lineage (SCL) [6, 7]. These stem-like tumor cells can drive tumor progression and relapse, contribute to varying therapeutic responses among patients, and consequently complicate treatment options [8]. On the other hand, the aberrant signals retained from NCSCs or SCL precursors (SCPs) present potential targets for possible alternative management strategies. Recent research on PNs and other neural crest-derived tumors has emphasized the targeting of highly expressed ERBB3, proposing novel treatment approaches [9, 10].

ERBB family consists of a group of receptor tyrosine kinases (RTKs) that play crucial roles in various cellular functions, such as growth, differentiation, and survival. This family includes four primary members: ERBB1 (also known as EGFR), ERBB2, ERBB3, and ERBB4. Each of these receptors can form homo-or heterodimers that significantly influence their signaling capabilities and functional implications. The dysregulation of the ERBB family component is a common feature in various tumor types, including PNs and MPNSTs [11, 12], and the intrinsic heterogeneity can be reflected by the differential dependency on the ERBB family receptors [13]. ERBB3 emerges as a crucial player in SCL development upon neuron regulations, such as NRG1, through its unique ability to form heterodimers with other ERBB family members, particularly ERBB2 [14, 15]. Our single cell RNA sequencing (scRNAseq) analysis reveals high *ERBB3* expression in the stem-like tumor population in human atypical neurofibromatous neoplasms of uncertain biologic potential (ANNUBP), the transition stage from PN to MPNST, and mouse MPNSTs [6]. In various cancers, including breast cancer and glioblastoma, ERBB3 dependency is apparent, often operating in conjunction with other kinases to regulate critical survival and proliferation signaling pathways [16]. Specifically, tyrosine-1289 (Tyr1289) phosphorylation of ERBB3 directly recruits PI3K, making ERBB3 a key activator of PI3K/AKT signaling [16, 17]. Other RTKs, such as EGFR, were also reported to regulate NF1-associated MPNSTs [18].

Heat shock protein 90 (HSP90) is a family of well-known molecular chaperones that facilitate the maturation and stable conformation of client proteins, including p53, serine/threonine kinases, receptor/non-receptor kinases, and steroid hormone receptors [19-22]. The HSP90 family is comprised of several isoforms, including HSP90α, HSP90β and GRP94, each exhibiting unique expression patterns and regulatory mechanisms within different cellular contexts [23]. The roles of the HSP90 family are even more prominent in the stem cells in balancing pluripotency and development [24, 25]. The use of HSP90 inhibitors (HSP90i) also has garnered attention in antitumor applications. HSP90i were reported to selectively eradicate lymphoma stem cells via HIF2α mediated hypoxia response [19] and effectively target glioblastoma cancer stem cells by downregulating the phosphorylation of key client proteins such as EGFR, ERBB3, and AKT [26]. Furthermore, HSP90i have been shown to inhibit MPNSTs by inducing unfolded protein response-related endoplasmic reticulum stress, a process that can be potentiated through co-administration with Rapamycin [26, 27]. These studies emphasize the crucial roles of HSP90 and suggest promising HSP90-targeted strategies for various tumors and cancer stem cells.

Our recent study on PN drug screen mining suggested that 1) there are conserved gene regulation networks among the PN cell lines and primary PN that reflect the conserved stem-like molecular properties, 2) several HSP90i show a propensity to preferentially inhibit PN cell lines with stem-like properties compared to others [28], and therefore, we propose that HSP90 inhibition can be essential for targeting the stem-like tumor cells with high ERBB3 in PNs. A Phase I clinical trial of the HSP90i SNX-5422 demonstrated good tolerance and antitumor effects when used in combination therapy for non-small cell lung cancer [29]. Ganetespib, another HSP90i, was also reported to have inhibitory effects in preclinical thyroid cancer models [30]. SNX-2112, a potent small molecule HSP90i, has been evaluated in hematological malignancies, including multiple myeloma and chronic lymphocytic leukemia. SNX-2112 has been shown to bind to various isoforms of the HSP90 proteins with differential affinities at the ATP-binding site, leading to the inhibition of these proteins and consequent loss of their chaperone activity [31, 32]. Clinical trials concerning SNX-2112 generally showcase its effectiveness in inducing apoptosis in multiple myeloma cells while impacting critical signaling pathways mediated by AKT and ERK, which are essential for cell identity and survival in cancer cells [33, 34]. Notably, a study revealed that SNX-2112 outperformed other conventional HSP90i, like 17-AAG, in various cancer cell lines [33].

In this study, we assess SNX-2112 and reveal its inhibitory effect in PNs, focusing on targeting PN cells with high *ERBB3* expression in both human and murine PN models. The fact that SNX-2112 decreases total and phosphorylated forms of ERBB3 and the downstream AKT activation demonstrated that this signaling can serve as a potential mechanism for its antitumor effects.

## Material and Methods

### PN transcriptome for gene enrichment analysis

The bulk sequencing data for PN cell lines and corresponding primary tissues were obtained from the NFdata portal via syn22351884. The analysis was conducted in RStudio with R version 3.6. Heterogenous tumor signatures from Schwann lineage tumor clusters CL0, CL1 and CL2 were collected from a prior study [6]. The GSVA R package was employed to compute the gene set enrichment score using default parameters [35]. The pheatmap R package was utilized to generate a heatmap that incorporates clinical information retrieved from the NFdata portal [36].

### Single cell analysis

The mouse scRNAseq data were downloaded via GSE181985 from Gene Expression Omnibus. The original data were collected from the 7-month-old pNF from *DhhCreNF1*^*f/f*^ mouse model. The data were processed by the default pipeline with R package Seurat. The doublets were removed, and the three different data sets were integrated by R package Harmony.

### Cell viability assay

CellTiter-Glow® assay was used to determine the effect of SNX-2112 on the cell viability according to the manufacturer’s instructions (REF G755B/G756B; Promega, USA). Briefly, exponentially growing cells were seeded in 96-well plates (5,000 cells/well) and allowed to attach overnight. The complete growth medium was then replaced with a DMEM-based medium containing increasing concentrations of SNX-2112 (0.04–0.32 μM) and incubated for 48 hours. After incubation, the plate was equilibrated to room temperature, and 20 μL of CellTiter-Glo reagent was added. The plate was placed on a shaker for 2-3 mins followed by incubation with the assay reagent for 7-8 mins at room temperature. The absorbance was then measured at 573 nm using SpectraMax® i3X microplate reader (software: SoftMax Pro 7.0.3), Molecular Devices, USA. The cell viability was normalized to the untreated control. Three independent replicates of the cell viability assay were performed to assess the effect of SNX-2112 on the viability of the treated cells.

### Western blot analysis and antibodies

After drug treatment, the cells were washed thrice with warm DPBS followed by lysis induced by cold Pierce™ RIPA buffer (REF 89901; Thermofisher, USA) containing protease inhibitor to a final concentration of 1% (v/v) and phosphatase inhibitor cocktail tablets (REF 04906837001, Roche, Germany). The protein content was quantified with Pierce™ BCA Protein Assay kit (REF 23225, Thermofisher, USA), according to the manufacturer’s instructions. Appropriate volumes of the extracted proteins were denatured, and comparable volumes of proteins were loaded on 7.5% Mini-PROTEAN^®^ TGX™ precast gels from Bio-Rad Laboratories Inc., USA, (Cat# 45568024) for separation, followed by transferring to a low-fluorescence PVDF blotting membrane (Cat. # 1620264; Bio-Rad Laboratories Inc., USA). The protein expression level was detected by overnight incubation at 4°C with primary antibodies for EGFR (Cat. # 4267), p-EFGR (Tyr1068; Cat. # 3777), ERBB2 (Cat. # 4290), ERBB3 (Cat. # 12708), p-ERBB3 (Tyr1289; Cat. # 2842), AKT (Cat. # 4691), p-AKT (Ser473; Cat. # 4060), and β-Actin (sc-47778; Santa Cruz Biotechnology, USA). Subsequently, the membranes were incubated with corresponding secondary antibodies at room temperature for up to 2 h. Restore™ Western Blot Stripping Buffer (REF 21059, Thermofisher, USA) was used to strip the membrane of the previous antibodies for further treatments when necessary. The proteins of interest were visualized with iBRight™ FL1000, Invitrogen by Thermofisher Scientific, USA (software version 1.8.2). The intensity of the detected proteins in images was quantified using Image J software (version V1.54p). After densitometric analysis, each protein band was normalized to respective β-actin and then to the untreated control. The experiments were replicated thrice for reproducibility and statistical reliability.

### In vivo experiment of SNX-2112

pNF-bearing dorsal root ganglia (DRG) isolated from adult *DhhCre;Nf1*^*f/f*^;*Luc* mice were cultured and injected into the sciatic nerve of nude mice. Following tumor formation, harvested tumors were dissociated and 5×10^5^ cells were transplanted into the sciatic nerve of 6-week-old nude mice. After confirming tumor engraftment using in vivo imaging system (IVIS) that detected luciferase activity, the mice were randomly assigned into experimental and control groups, with each group comprising of 5 animals. The experimental group received SNX-2112 at a dosage of 30 mg/kg every 48 hours intraperitoneally (I.P.), while the control group received vehicle only composed of 45% normal saline, 40% polyethylene glycol 400, 5% Tween 20, and 10% dimethyl sulfoxide (DMSO). Tumor growth was monitored twice a week by IVIS to assess luciferase signal intensity over a two-week period. At the study endpoint, all mice were euthanized, and tumors were harvested for subsequent tumor volume analysis. The animal handling and treatment complied with the mouse protocol #7457, which was approved by the IACUC of Medical College of Wisconsin.

### Statistical analysis

Graphs and statistical analyses were generated using GraphPad Prism, version 10.4.0 (GraphPad Software, San Diego, CA, USA). The data were presented as mean ± SD of three independent replicates. Statistical significance was determined using one-way ANNOVA to compare treatment groups to untreated control, two-way ANNOVA to determine the effect of the drug and duration of treatment, and student’s t-test for two-group comparisons. A p-value less than or equal to 0.05 was considered statistically significant.

## Results

### PN cell lines retain molecular features of PN intrinsic heterogeneity

The primary NF1-associated tumor cells are hard to culture because of their slow-growing rate and senescent phenotypes in vitro [37]. PN cell lines provide practical models to study this rare tumor in vitro, however, it is pivotal to understand their molecular differences, reflecting possible intrinsic tumor heterogeneity within the PN [38]. Dr. Wallace’s (Peggy) team generated a series of PN cell lines overcoming the growth bottleneck by overexpressing mouse CDK4 and human telomerase reverse transcriptase [39]. To understand the molecular features, we characterized the transcriptome of PN cell lines, adjacent Schwann cell lines, and corresponding primary tumors from the patients (Figure 1A). We applied the gene set enrichment analysis (GSEA) using gene signatures from the scRNAseq of a human ANNUBP. The three gene sets represented the intrinsic heterogeneity and different magnitudes of stemness. CL0.sig and CL1.sig enrich SCL precursors (SCPs) signatures, whereas CL2.sig has the genes involved in differentiated and senescent Schwann cells with the least stem potential (CL0.sig > CL1.sig > CL2.sig) [6]. The GESA score heatmap shows that while the majority of primary PN samples have higher stemness scores (CL0.sig or CL1.sig), the corresponding cell lines have much lower CL0.sig scores but higher CL2.sig scores. These interesting contrasts indicate that the primary PNs, such as p95.11bC, p97.4, and p04.4, have a significant number of the stem-like population, and pherhaps via the technical processing and selection during the immortalization, the stem-like signatures may be stochastically maintained in the tumor cell lines, such as ipNF95.11bC (*NF1*^−/−^) and ipNF05.5 Mixed Clone (*NF1*^−/−^). NCSC signature genes, such as *ERBB3* and *SOX10*, consistently decrease from the primary PN to the corresponding cell lines (Figure 1B&C). This systematic transcriptome comparison enabled us to understand and exploit different cell lines as surrogates for heterogeneous PN populations and evaluate the treatment strategy targeting the ERBB3^high^ molecular feature.

**Fig 1.**
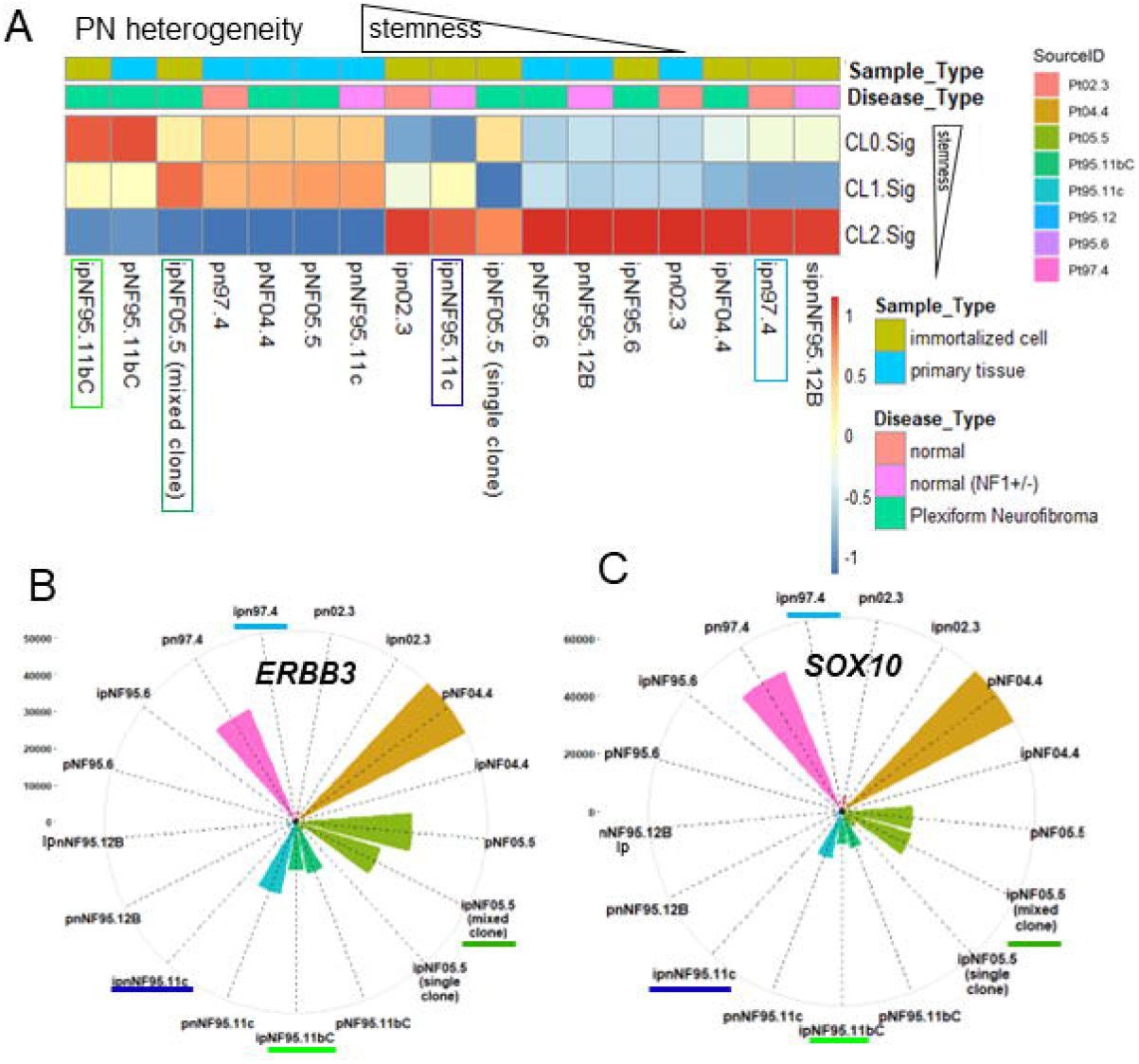

### PN cell lines demonstrate different ERBB family preference

To further evaluate the diversity of the ERBB family, we determined the ERBB3 and EGFR protein levels in the PN cell lines. ipNF95.11bC and ipNF05.5 Mixed Clone demonstrate the elevated total ERBB3 level, while ipnNF95.11c and ipn97.4 show relative high EGFR (Figure 2A&B). ERBB2, the most common partner to form heterodimers with ERBB3, was found to be highly expressed on the ipNF95.11bC and ipNF05.5 Mixed Clone cells, which is consistent with ERBB3 pattern, as shown in Figure 2C. The co-existence of ERBB family members may reflect the dynamics of tumor cells to adapt to the environment. In the context of glioblastoma, ERBB3 has been implicated in promoting resistance to EGFR-targeted therapies, where cells expressing both EGFR and ERBB3 can activate downstream PI3K signaling regardless of the functional status of EGFR. ipNF95.11bC and ipNF05.5 mixed clone showed significantly lower expression of EGFR and higher expression of ERBB3 and ERBB2 when compared to ipnNF95.11c and ipn97.4 (Figure 2D, E, and F), suggesting the differential ERBB member dependency in the PN cell lines. These data validate the molecular characterization in Figure 1 and further facilitate us in testing the HSP90i on the PN cells with ERBB3^high^ molecular feature.

**Fig 2.**
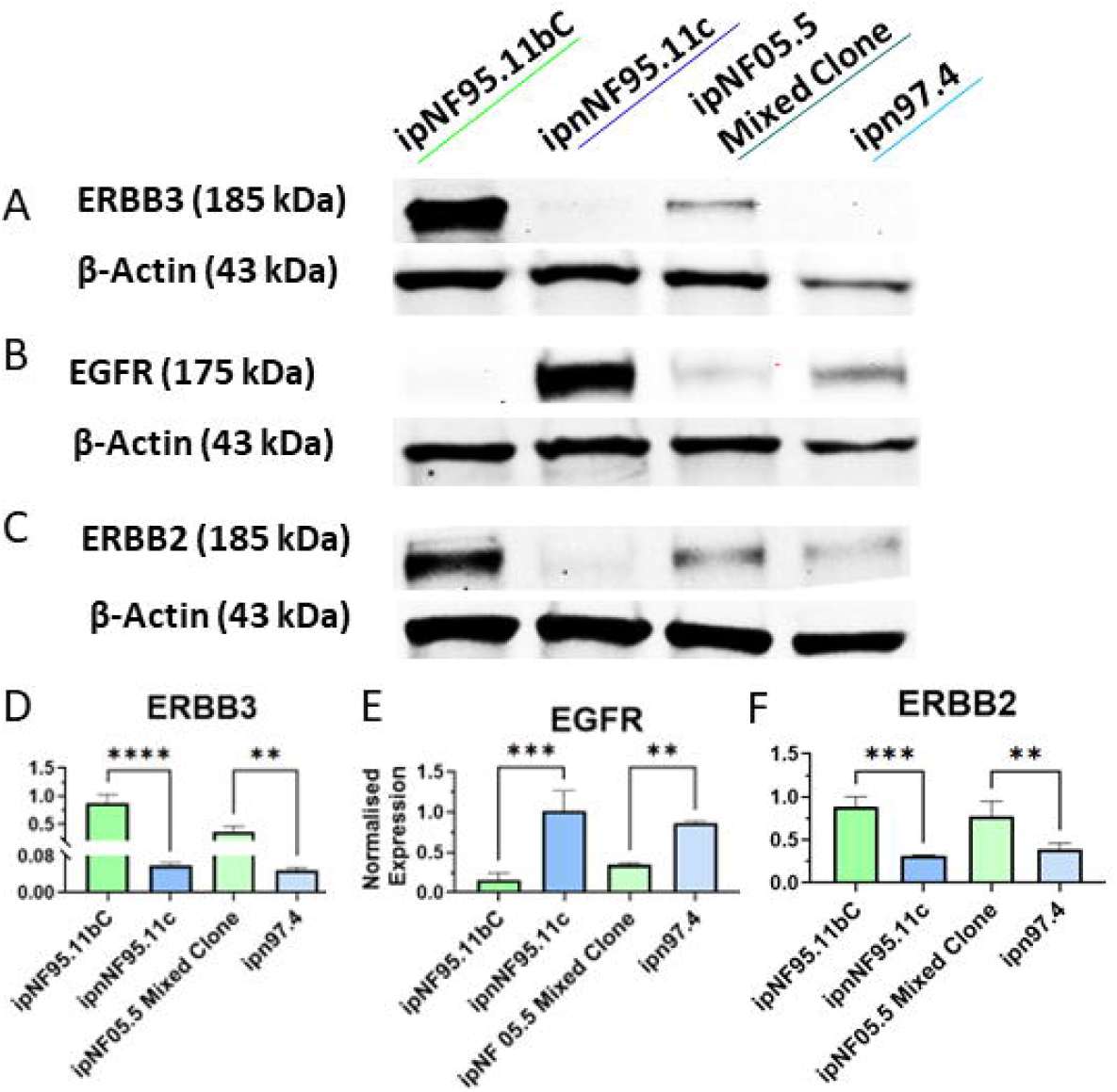

### Differential drug responses of the PN lines to SNX-2112

We analyzed the drug screen data published in Marc Ferrer’s study, in which the MIPE 4.0 drug library was applied to multiple PN cell lines to discover drug candidates [40]. The drug response heatmap demonstrated that the ERBB3^high^ cell lines, ipNF95.11bC and ipNF05.5 Mixed Clone, are more sensitive to the HSP90i than ipnNF95.11c (Figure 3A). We validated the drug responses using one of the HSP90i, SNX-2112, on four Schwann cell lines (Figure 3B). The CellTiter-Glow® assay showed a concentration-dependent effect of SNX-2112 on the viability of the tested cell lines. The differential responses between the ERBB3^high^ (green lines) and ERBB3^low^ (blue lines) were confirmed. Additional HSP90i, including SNX-5422, Retaspimycin and Geldanamycin, were tested on these cell lines, and significant vulnerability was observed in the ERBB3^high^ cell lines (Supplementary Figure 1). The results strongly demonstrated the vulnerability of ERBB3^high^ PN cell lines to HSP90i.

**Fig 3.**
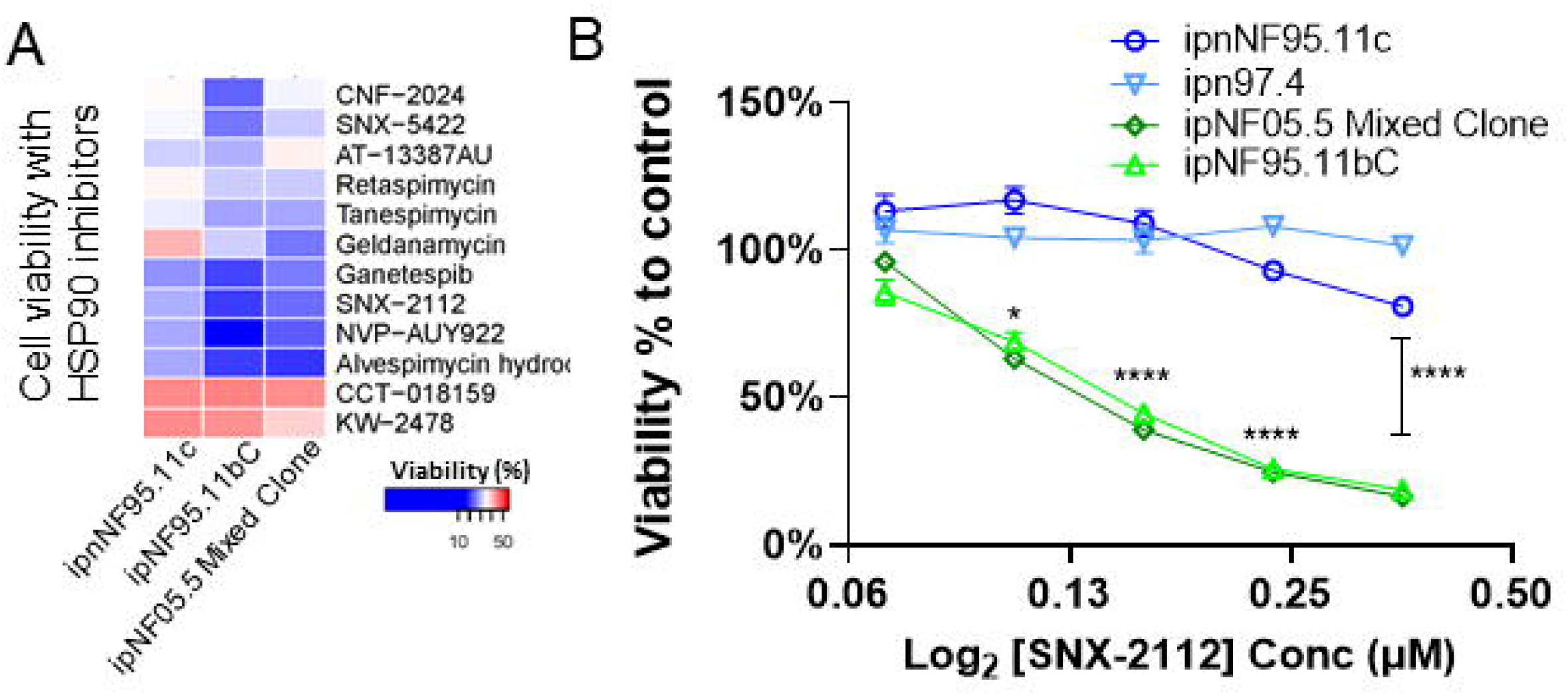

### SNX-2112 decreases the total ERBB3 and p-ERBB3

To assess the ERBB3-related inhibitory effects from SNX-2112, western blots were performed to determine ERBB3 expression levels in the cell lines following SNX-2112 treatment with or without NRG1 stimulation. Notably, the dose-dependent decrease in total ERBB3 is more pronounced in ipNF95.11bC and ipNF05.5 Mixed Clone than in the control cell lines (Figure 4A&B). The quantification of the total ERBB3 of the cell lines is in Supplementary Figure 2A&B. Additionally, activated p-ERBB3(Tyr1289) was significantly reduced in the ipNF95.11bC and ipNF05.5 Mixed Clone compared to ipnNF95.11c and ipn97.4 following SNX-2112 treatment even with NRG1 stimulation (Figure 4C&D). The quantification of p-ERBB3(Tyr1289) of the cell lines is in Supplementary Figure 2C&D. Phosphorylated ERBB3 antibodies at Tyr1187 and Tyr1249 were evaluated, but no bands were detected (data not shown). These findings demonstrated a differential impact of SNX-2112 on cells with ERBB^high^ versus other Schwann cells, which accounts for more significant cytotoxic effects observed in ipNF95.11bC and ipNF05.5 Mixed Clone cells relative to ipnNF95.11c and ipn97.4.

**Fig 4.**
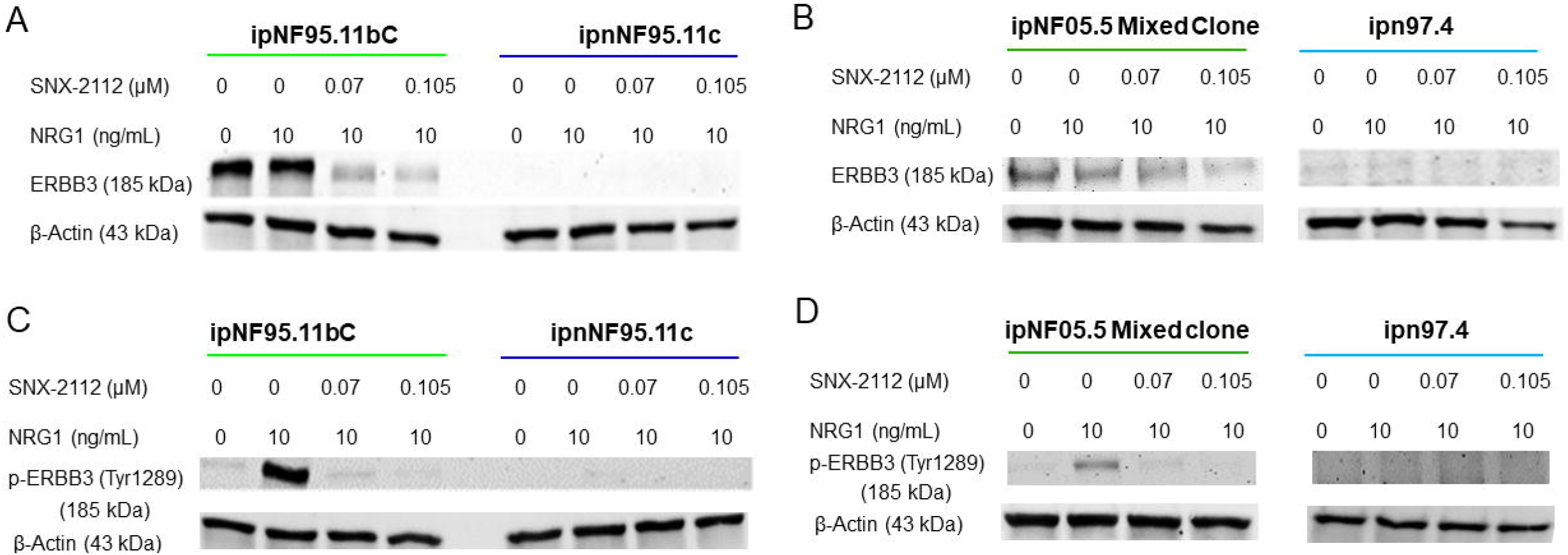

### SNX-2112 inhibits total AKT and p-AKT

The aberrant AKT signaling contributes to enhanced tumor cell growth and survival under stress conditions, such as chemotherapy and radiotherapy, thereby increasing treatment resistance [41]. Activation of ERBB3 has been reported to enhance the recruitment of the p85 subunit of the PI3K to the plasma membrane, subsequently promoting AKT activation, which takes place downstream from the binding of NRG1 to ERBB3 during early SCL development [42]. AKT signaling can also involve SCL differentiation and the thickness of the myelin sheath [31, 43]. NF1-associated tumors were reported to have activated PI3K/AKT signaling [44, 45]. The ERBB3-dependent regulatory mechanism can be exploited from the SCL precursors, and the constitutive activation of the AKT pathway can be essential for stem-like cell-driven tumorigenesis [46, 47]. To explore the possible downstream inhibitory mechanisms of SNX-2112, we performed a western blot to evaluate the total AKT and p-AKT (Ser473). As shown in Figure 5 A and B, a significant dose-dependent decrease of total AKT upon the SNX-2112 treatment was observed (p<0.05). The western blot quantification of the total AKT and p-AKT (Ser473) were demonstrated in Supplementary Figure 3. More interestingly, SNX-2112 significantly decreased p-AKT(Ser473) in ipNF95.11bC and ipNF05.5 Mixed Clone lines, and the activation of AKT was not detected in the control cell lines (Figure 5C&D). This suggests that ERBB3^high^ cell lines have higher basal level of p-AKT(Ser473), which can be further activated via the NRG1-ERBB3 axis in these cells. Vulnerability due to decreased p-AKT (Ser473) in the ERBB3^high^ cell lines may explain the dose-dependent inhibition of SNX-2112 in Figure 3B, while the viability of ERBB3^low^ cell lines remained largely unaffected, which highlights the possible mechanism of susceptibility of ERBB3^high^ cells to SNX-2112.

**Fig 5.**
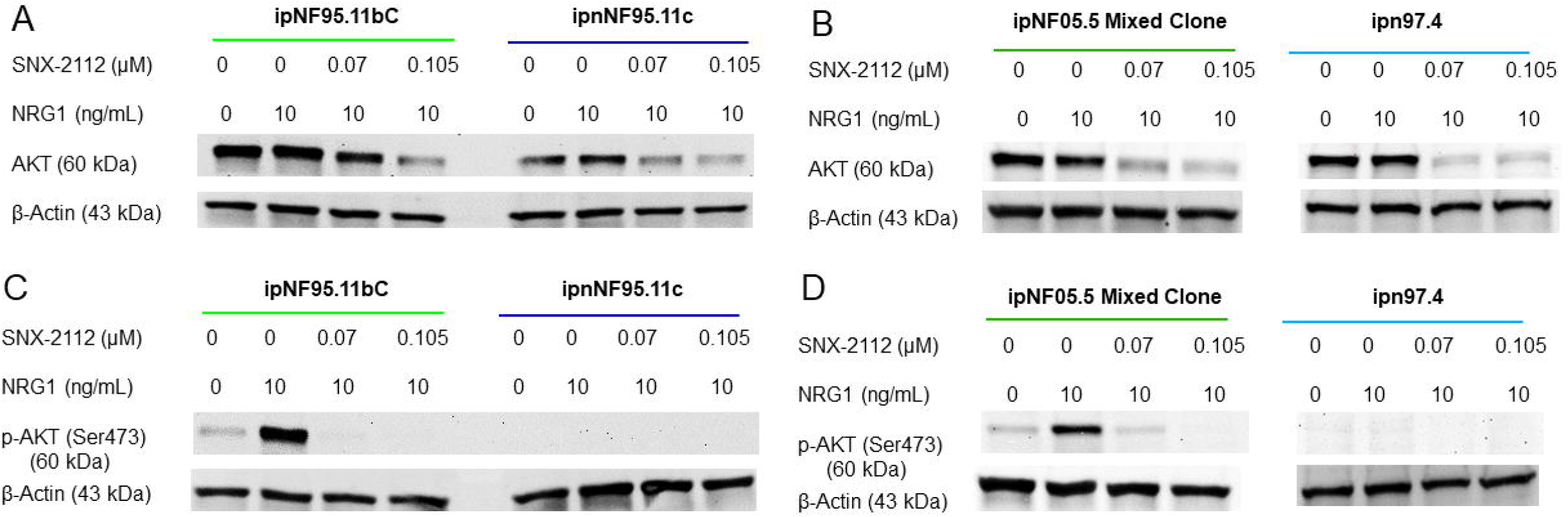

### SNX-2112 inhibits PNs from *DhhCre;NF1*^*f/f*^ models in vivo

The ERBB3^high^ tumor cell population was characterized in both human patients and multiple mouse models, including *DhhCre;Nf1*^*f/f*^ [6, 9, 48]. The elevated *Erbb3* expression was further confirmed by scRNAseq analysis using data from 7-month-old *DhhCre;Nf1*^*f/f*^ PN at paraspinal dorsal root ganglia and wildtype control (Supplementary Figure 4A). The *Erbb3* expression was highly enriched in the stem-like SCL in the 7-month tumor compared to the control (Supplementary Figure 4B&C). To test the inhibitory effects of SNX-2112 on PNs in vivo, cells dissociated from *DhhCre;Nf1*^*f/f*^;*Luc* were injected into the sciatic nerves of 10 nude mice to generate the PN transplantations. A total of 5×10^5^ PN tumor cells were injected for each mouse, and the mice were randomly split into two cohorts. Drug treatment began four days post-transplantation, with either SNX-2112 (30 mg/kg) or vehicle being injected intraperitoneally to the mice every two days (Figure 6A). Bioluminescence was used to track the tumor growth twice per week (Figure 6B). A significant difference in tumor progression was demonstrated between the two cohorts after the treatment (p<0.01). The representative bioluminescence images at different time points are shown in Supplementary Figure 5. At the endpoint, the tumors were dissected, and a significant difference in tumor volume from the two cohorts was shown in Figure 6C (p<0.05). The comparison between two cohorts of PN tumors was demonstrated in Figure 6D, which shows the significant tumor inhibitory effects of the SNX-2112 treatment. No significant toxicity was observed and reflected by weight loss in the cohorts (data not shown).

**Fig 6.**
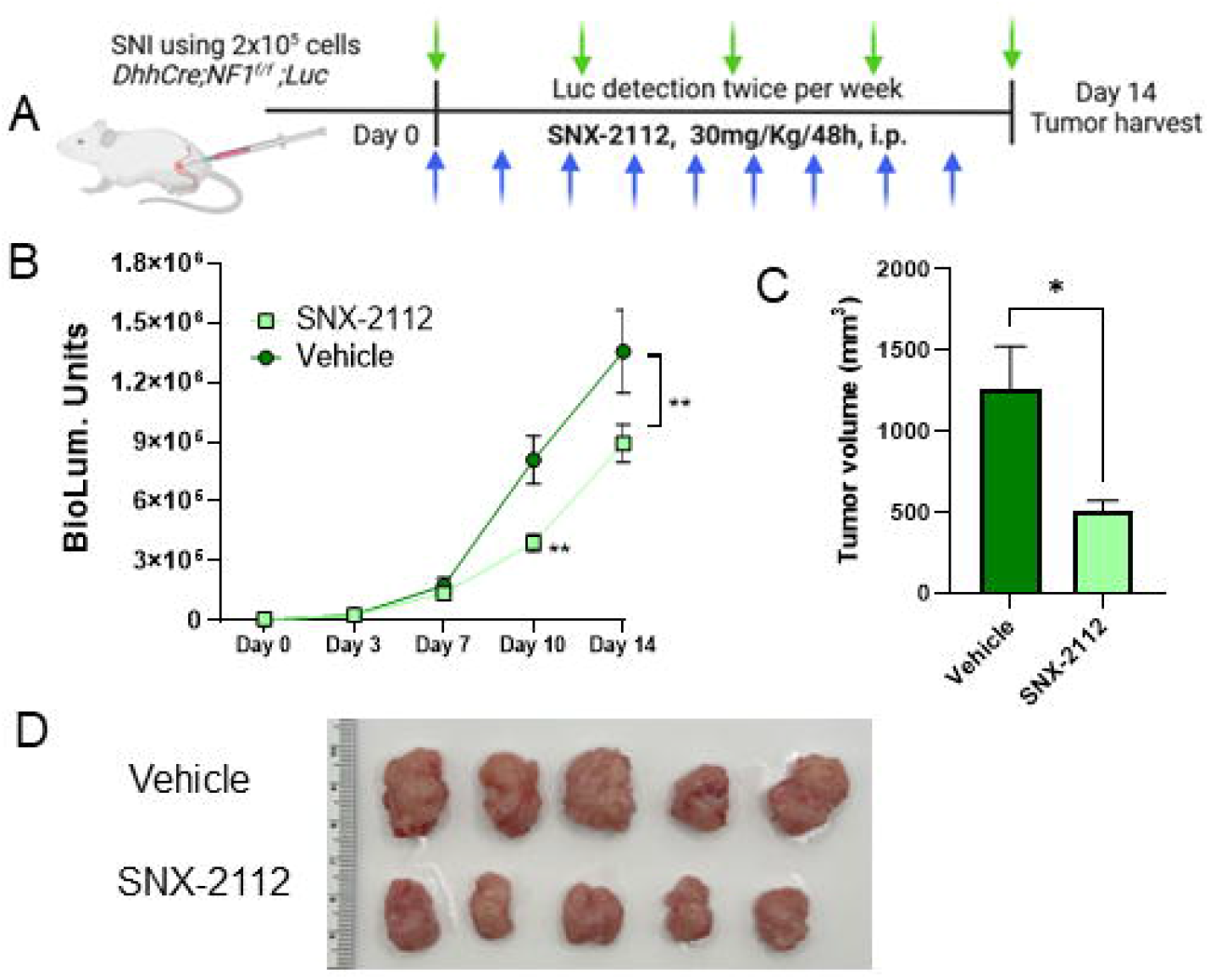

## Discussion

As a type of benign tumor, PNs are challenging to culture *in vitro*, which makes the immortalized cell lines practical tools for exploring new treatments, especially in drug screens. However, the intrinsic tumor heterogeneity may be partially reflected by the immortalized cell lines established from a specific specimen, depending on the complex processes of biopsies and clone selection. Meanwhile, the establishment of numerous cell lines may capture the PN heterogeneity through a stochastic process, which can be determined by transcriptome variations among the cell lines. Characterization of the molecular features of these cell lines is essential to understanding the differential drug responses and enabling the researchers to study the drug-targeted molecular features resembling the heterogeneous tumors [49, 50]. In this study, we compared the transcriptomes from matched PNs and SCL cell lines to uncover a new angle to take advantage of molecular features of cell lines, such as ipNF95.11bC for high ERBB3 expression, and use these cells to study the drug vulnerability of ERBB3^high^ PNs. Our characterization demonstrated differential dependencies on ERBB family members in the immortalized SCL cell lines, which may explain the variations among of the responses in the drug screen [40].

PN treatment presents significant clinical challenges, as these tumors often arise at a very young age and may exhibit varying degrees of response to MEK inhibitor therapies. The aberrantly high ERBB3 expression can serve as a targetable molecular feature of stem-like cells in PNs, and its regulation of the downstream AKT activation offers a mechanism through which the therapies can be developed. The findings in this study support the use of HSP90i, specifically SNX-2112, as a treatment option for PNs. The in vivo efficacy of SNX-2112 in reducing tumor volume provides a promising outlook on the potential translation of HSP90 inhibition into the clinical practice on PNs.

By focusing on the stem-like features within PNs, our research elucidates a promising avenue for a therapeutic strategy that considers the unique properties of this population. More importantly, early intervention is essential in preventing tumor progression in pediatric patients. The stem-like tumor population with high expression of *ERBB3* is known as the driver in PN progression [6, 9, 16]. Targeting this population may significantly benefit pediatric patients. The downregulation of ERBB3 and the inhibition of downstream AKT signaling not only mechanistically explains tumor inhibition with the use of SNX-2112, but also may pave the way for more tailored therapeutic approaches at an early stage of PN.

Using HSP90i inhibitors represents a shift towards more targeted medicine, potentially alleviating symptoms and improving the quality of life for patients afflicted by this challenging condition. Our data supports the notion that effective treatment of PNs will benefit from the specific targeting of the pathways critical to the survival of stem-like tumor cells, thereby improving patient outcomes. The present study further elucidates the role of intrinsic heterogeneity in PNs and supports the deployment of HSP90i as a strategic approach to managing these tumors. Future investigations should explore the long-term effects of such treatments and assess combination strategies to enhance therapeutic efficacy against NF1-associated malignancies.

## Supporting information

Supplementary Figure 1 related to Figure 3

Supplementary Figure 2 related to Figure 4

Supplementary Figure 3 related to Figure 5

Supplementary Figure 4 related to Figure 6

Supplementary Figure 5 related to Figure 6

## Acknowledgments

We acknowledge Dr. Wallace’s generosity in providing the PN cell lines and inspiration consulting on the PN cell phenotypes.

## Author contributions

S.K. and D.S. for conceptualization and original writing; S.K., O.A. and B.B for data curation and formal analysis; D.S. for Funding acquisition; X.V. and D.A. for Resources; K.N. for Visualization; D.S. for Supervision; L.S. and H.Z. for editing and revision

