## Supplementary figures and images for "SNX-2112 inhibits the ERBB3^high^ tumor population in Plexiform Neurofibromas"

### Supplementary Figure 1 related to Figure 3

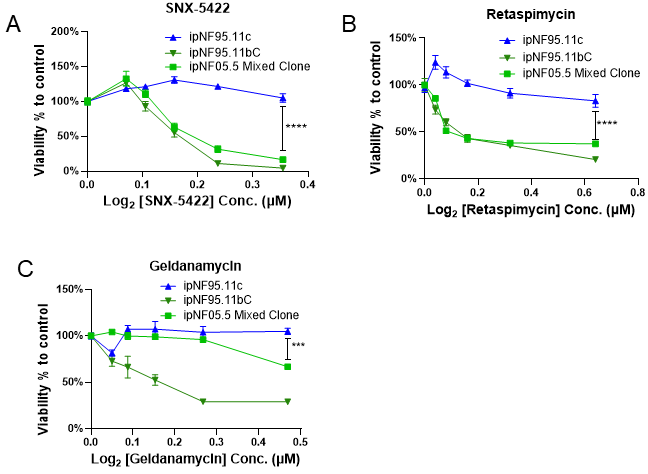

### Supplementary Figure 2 related to Figure 4

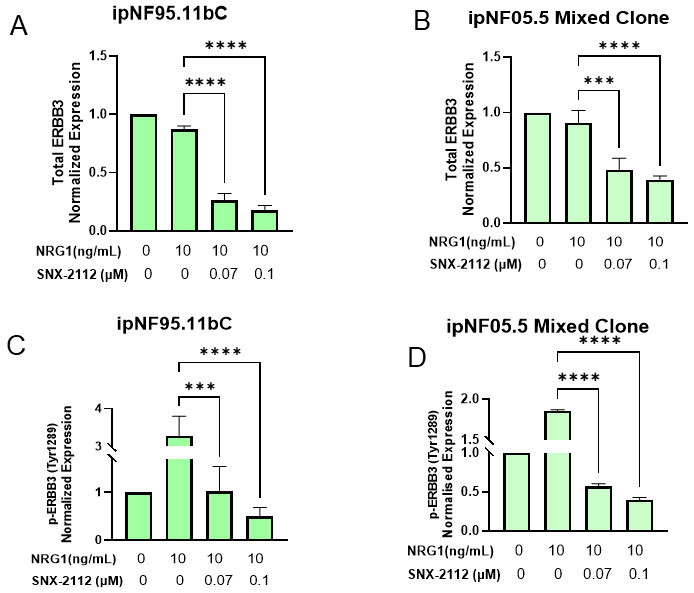

### Supplementary Figure 3 related to Figure 5

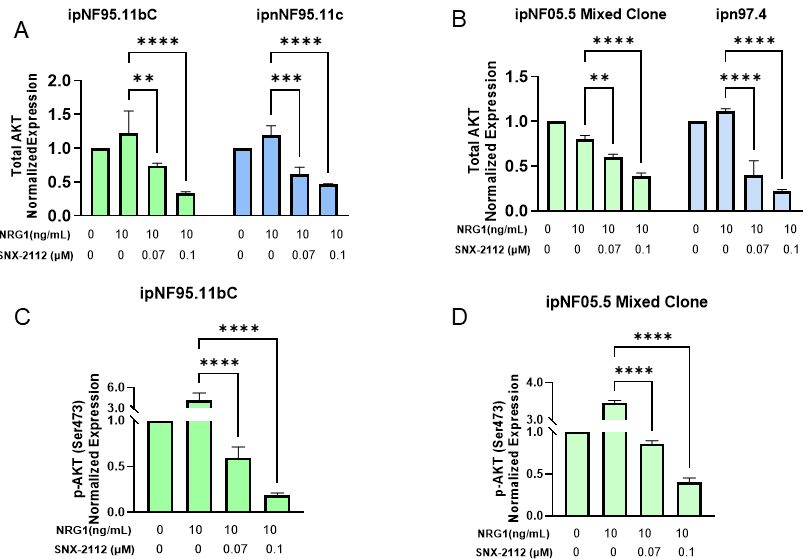

### Supplementary Figure 4 related to Figure 6

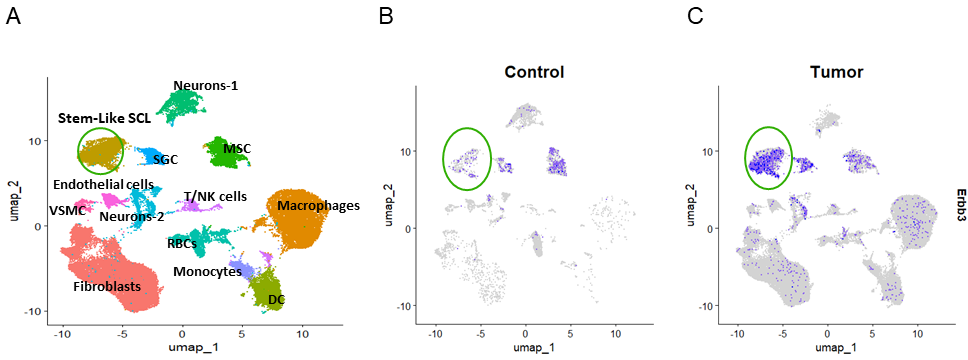

### Supplementary Figure 5 related to Figure 6

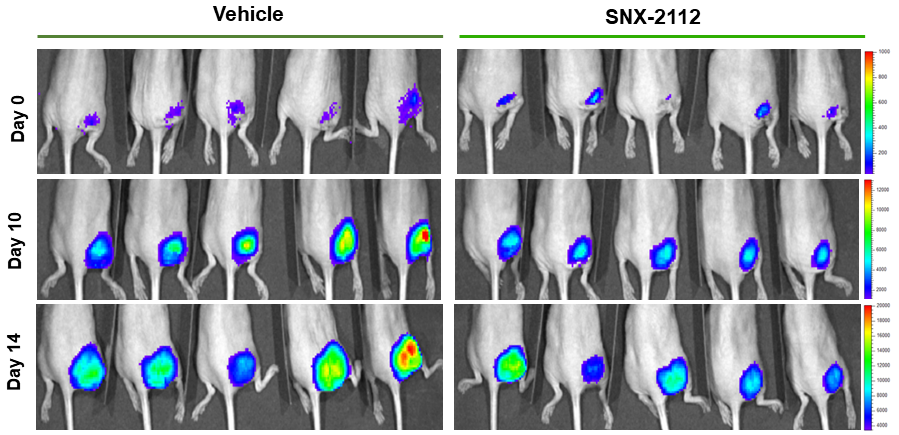
